# Zygotic pioneer factor activity of Odd-paired/Zic is necessary for establishing the *Drosophila* Segmentation Network

**DOI:** 10.1101/852707

**Authors:** Isabella V. Soluri, Lauren M. Zumerling, Omar A. Payan Parra, Eleanor G. Clark, Shelby A. Blythe

**Affiliations:** Department of Molecular Biosciences, Northwestern University, Evanston, IL; Program in Interdisciplinary Biological Sciences, Northwestern University, Evanston, IL; Department of Neurobiology, Northwestern University, Evanston, IL

## Abstract

Because regulatory networks of transcription factors drive embryonic patterning, it is possible that chromatin accessibility states impact how networks interact with information encoded in DNA. To determine the interplay between chromatin states and regulatory network function, we performed ATAC seq on *Drosophila* embryos over the period spanning the establishment of the segmentation network, from zygotic genome activation to gastrulation. Chromatin accessibility states are dynamic over this period, and establishment of the segmentation network requires maturation of the ground chromatin state. Elimination of all maternal patterning information allows identification of patterning-dependent and -independent dynamic chromatin regions. A significant proportion of patterning-dependent accessibility stems from pioneer activity of the pair-rule factor Odd-paired (*opa*). While *opa* is necessary to drive late opening of segmentation network cis-regulatory elements, competence for *opa* to pioneer is regulated over time. These results indicate that dynamic systems for chromatin regulation directly impact the interpretation of embryonic patterning information.

## Introduction

Embryonic patterning systems direct a set of initially uncommitted pluripotent cells to differentiate into a variety of cell types and complex tissues. Over developmental time spans, regulatory networks of transcription factors drive the acquisition of unique cell fates by integrating patterning information and determining the set of genes to be activated or repressed in response to developmental cues [1]. The critical nodes of these regulatory networks are cis-regulatory modules (CRMs) where transcription factors bind in order to enhance or silence target gene activity. However, additional epigenetic determinants such as the organization of chromatin structure likely influence how genomic information is accessed by regulatory networks. For instance, because nucleosome positioning can hinder transcription factor-DNA interactions [2, 3] chromatin effectively serves as a filter either to highlight or obscure regulatory information encoded in DNA. But embryonic chromatin states themselves are dynamic [4–9]. The mechanisms that drive developmental progression can also trigger remodeling of chromatin accessibility patterns on both large and small scales, thereby changing over time what genetic information is available to gene regulatory systems. While in many cases we have near comprehensive understanding of both the genetic components of certain developmental networks and the critical CRMs whereby these components interact, much less is known about how chromatin accessibility states constrain network function and how mechanisms for controlling chromatin accessibility are systematically woven into the developmental program.

In the case of *Drosophila melanogaster*, decades of investigation into the mechanisms of development have exhaustively identified the critical patterning cues and transcription factors that drive early cell fate specification and differentiation of select developmental lineages. Patterning is initiated by four distinct maternal pathways that alone are sufficient to initiate zygotic regulatory networks that specify all of the primary cell identities that arise along the major embryonic axes [10–17]. At the outset of patterning, nuclei have what can be considered a ‘ground state’ of chromatin structure that contains the initial set of accessible CRMs and promoters that will define the first regulatory network interactions [7]. The ground state effectively provides a baseline for determining the influence of epigenetic mechanisms of gene regulation on developmental processes. The early *Drosophila* embryo therefore provides an ideal starting point to observe both how regulatory networks are constrained by chromatin states, and how these states evolve as a function of progression through the developmental program.

Before embryos can respond zygotically to maternal patterning cues, they must first undergo a series of 13 rapid, synchronous mitotic divisions that serve to amplify the single nucleus formed after fertilization into a set of ∼6000 largely uncommitted, pluripotent cells [18, 19]. These mitotic divisions occur in a state of general transcriptional quiescence that effectively prevents nuclei from responding prematurely to regulatory stimuli [20–23]. The shift from the initial proliferative phase to later periods of differentiation comes at a major developmental milestone termed the midblastula transition (MBT) during which the zygotic genome activates, and cells become competent to respond to maternal patterning information [24]. A major component of zygotic genome activation (ZGA) is the establishment of the chromatin ground state by a subset of transcription factors known as pioneer factors that gain access to closed or nucleosome-associated DNA and direct the establishment of short tracts of open and accessible chromatin [25–31]. Because maternal pioneer factors such as Zelda are expressed uniformly in all cells [25], it is inferred that the initial ground state is common to all cells of the embryo at ZGA. However, maternal patterning systems can directly influence chromatin accessibility states at ZGA [32], and because their activities are by definition spatially restricted, this gives rise to heterogeneous embryonic chromatin states. Indeed, ATAC-seq measurements of chromatin accessibility in subdivided post-ZGA embryos, sampled either as posterior and anterior halves [33], sorted cells purified from restricted positions along the anterior-posterior (AP) axis [34], or as single cells [8] identify spatial heterogeneities in accessibility patterns that begin to emerge shortly after embryos undergo ZGA and initiate patterning. These observations raise the question of what mechanisms drive the reshaping of the chromatin landscape following ZGA.

In this study, we have investigated how chromatin accessibility states change following ZGA and to what extent these changes are dependent on the mechanisms of embryonic patterning. We find that the ZGA chromatin state must continue to evolve in order to support the establishment of accessible CRMs within the regulatory network that confers embryonic segmental identities. By measuring changes in chromatin accessibility over the one-hour period between ZGA and gastrulation comparing wild-type and mutant embryos in which all graded maternal inputs to patterning are either eliminated or flattened, we define sites that display dynamic regulation of accessibility downstream of either localized pattern-dependent or global patterning-independent cues. We find that although maternal patterning systems are limited in their ability to influence directly chromatin accessibility states, distinct downstream components of zygotic gene regulatory networks make major contributions to patterning-dependent alterations of the chromatin accessibility landscape. We focus on the characterization of one such factor, Odd-paired (Opa), which we demonstrate is both necessary and sufficient to pioneer open chromatin states for a set of cis-regulatory elements critical for the function of the embryonic segmentation network. These results highlight that individual components of gene regulatory networks may operate not simply to activate or repress target gene expression, but to dictate how and when network components interact by controlling dynamic cis-regulatory element accessibility.

## Results

### The ZGA Chromatin State is Insufficient to Support Embryonic Segmentation

To estimate the sufficiency of the ZGA chromatin state to support early embryonic development, we scored all known enhancers in the anterior-posterior (AP) segmentation network for chromatin accessibility. During ZGA, the first zygotic components of the segmentation network are transcribed in response to maternal patterning cues [35]. Over the course of nuclear cycle 14 (NC14) complex patterned gene expression from the gap, pair-rule, and segment polarity networks sequentially drive the conversion of graded maternal information into the unique segmental identities of cells across the AP axis [16, 36–39]. By entry into gastrulation, approximately 65 minutes into NC14, the initial activation of all three networks is complete. Because many of the CRMs required to generate the complex expression patterns of genes within the segmentation network are known, we used this system as a model to ask whether the ZGA chromatin state contains all the accessible cis-regulatory information required to complete this well-characterized developmental patterning task.

To evaluate chromatin accessibility states between ZGA and gastrulation, we collected single embryos aged either 12 or 72 minutes into NC14 and performed ATAC-seq. Mapped reads were assigned to peaks, which were subsequently cross-referenced against the Redfly database of previously characterized CRMs within the segmentation network [40]. These were then scored for accessibility either shortly after ZGA (NC14+12’) or one hour later at the onset of gastrulation (NC14+72’) (Figure 1A; see Materials and Methods).

**Figure 1:**
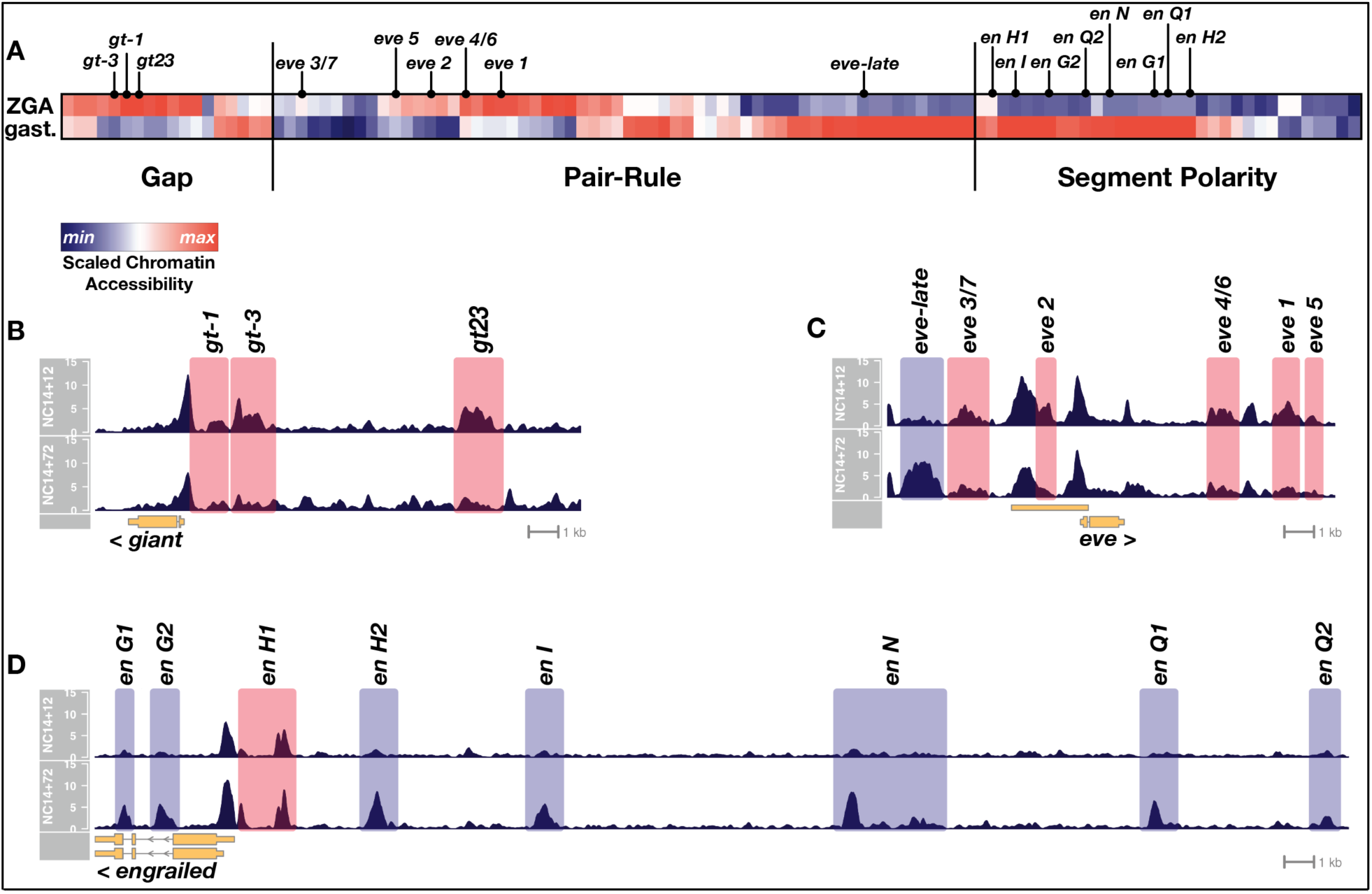
Chromatin accessibility at segmentation network enhancers is dynamic over the one-hour period between ZGA and gastrulation. A) Heatmap showing the scaled chromatin accessibility between ZGA (top row, NC14 + 12’) and gastrulation (bottom row, NC14 + 72’) over the complete set of known enhancers within the gap, pair-rule, and segment polarity gene regulatory networks. Tiers of the segmentation network are indicated as well as selected enhancers from the specific examples depicted in panels B-D. (B-D) chromatin accessibility at a representative gap (giant), pair-rule (eve), and segment polarity (engrailed) locus between ZGA (NC14+12, top) and gastrulation (NC14+72, bottom). Enhancers that are significantly more open by ZGA are highlighted in red, those that open late, by the onset of gastrulation are highlighted in blue. Loci are drawn to the same genomic (x-axis) scale, and ATAC accessibility is shown at the same y-axis scale (0-15 CPM) across all plots.

We find that CRMs within each tier of the segmentation network are not constitutively accessible and have distinct temporal chromatin accessibility profiles that correlate with the activity periods associated with these regulatory elements. All of the known early gap gene CRMs are open at ZGA and typically lose accessibility by the onset of gastrulation (Figure 1A, B). In contrast, pair-rule CRMs separate into two distinct temporal classes of chromatin accessibility. All of the early, stripe-specific CRMs within the pair-rule network are open at ZGA, whereas later, seven-stripe (or 14-stripe) specific CRMs generally lack open chromatin at ZGA, and gain accessibility by the onset of gastrulation (Figure 1A, C). The majority of the known segment polarity CRMs lack accessible chromatin at ZGA and undergo significant gains in accessibility by gastrulation (Figure 1A, D). Taken together, these results demonstrate that during the one-hour period between ZGA and gastrulation patterns of chromatin accessibility within segmentation network CRMs are dynamic, correlating with the early or late activity of gene expression patterns within the network. We conclude from this that the ZGA chromatin state contains insufficient accessible cis-regulatory information to sustain the function of the segmentation gene network. Chromatin accessibility patterns continue to evolve over the one-hour period between ZGA and gastrulation to support the later-acting components of the network, particularly the segment polarity and late pair-rule systems. This raises the possibility that the hierarchical networks that drive embryonic segmentation derive timing information from regulated chromatin accessibility. Notably, binding of pioneers [28, 41] implicated in the establishment of the initial ZGA chromatin state is low or absent at sites that gain accessibility late (Figure 1 Supplement 1), suggesting that additional, unrecognized regulatory mechanisms drive further changes in chromatin accessibility after ZGA.

### Identification of patterning-dependent and -independent changes in chromatin accessibility

The post-ZGA changes in chromatin accessibility could arise either uniformly within all cells of the embryo or could stem from the localized effects of developmental patterning systems. Previous investigations of chromatin accessibility states in post-ZGA embryos have demonstrated that shortly after ZGA, chromatin accessibility is largely homogeneous across most cells of the embryo [32, 33], but as development proceeds, spatially restricted, lineage-specific patterns of accessibility emerge as cell fates are specified and spatially restricted gene expression programs mature [8, 34]. One major exception to ZGA chromatin state homogeneity is a small cohort of enhancers with anteriorly restricted accessibility whose open state depends on the activity of the maternal patterning factor Bicoid (Bcd) [8, 32, 33]. Besides Bcd, it is unclear whether any other maternal patterning systems or downstream zygotic targets directly impact the chromatin accessibility state of the early embryo. We therefore measured the effect of eliminating all graded maternal inputs to embryonic development on the changes in chromatin accessibility we observe between ZGA and gastrulation.

By eliminating or flattening all graded maternal inputs to development, we force all cells of the embryo to develop along a single restricted trajectory. In situ hybridization of wild-type NC14 embryos for representative markers of anterior-posterior (AP) and dorsal-ventral (DV) specification reveals distinct, spatially restricted gene expression patterns along both axes (Figure 2A, left column). Four maternal systems provide the information sufficient to distinguish these positional identities along the AP and DV axes. The terminal extrema of the embryo are distinguished by localized Torso receptor tyrosine kinase antagonism of the medial determinant Capicua [13, 14]. Anterior fates are specified by the Bcd transcription factor gradient [10, 42]. Posterior identity is conferred by clearance of maternal Hunchback transcriptional repressor by graded expression of the Nanos translational repressor [15, 17, 43]. DV positions are specified by a gradient of the transcription factor Dorsal that functions downstream of a ventral-to-dorsal gradient of activated Toll receptor [44]. We generated embryos from mothers with null mutations in all terminal and AP systems (*bcd*, *osk*, *cic*, *tsl*) as well as a “lateralizing” allele of *Toll* (*Tl^RM9^*) that yields uniform, mid-level Dorsal activity across the entire DV axis [44] (see Discussion). All cells in these quintuple mutant embryos are positive for markers of lateral (*sog*) and posterior-terminal (*byn*) positional markers and do not express markers from other regions of the embryo (Figure 2A, right column). By reference to a previously reported single-cell RNA-seq dataset [45], dual *sog+ byn+* cells constitute a small fraction of posterior-lateral cells at this stage of development (Figure 2B) that are all fated to become posterior endoderm. We predicted that, compared with wild-type embryos, *bcd osk cic tsl Tl^RM9^* (hereafter, “mutant”) embryos would enrich for posterior endodermal chromatin states at the expense of all others, allowing for the unambiguous determination of sites that undergo patterning-dependent versus -independent changes in accessibility.

**Figure 2:**
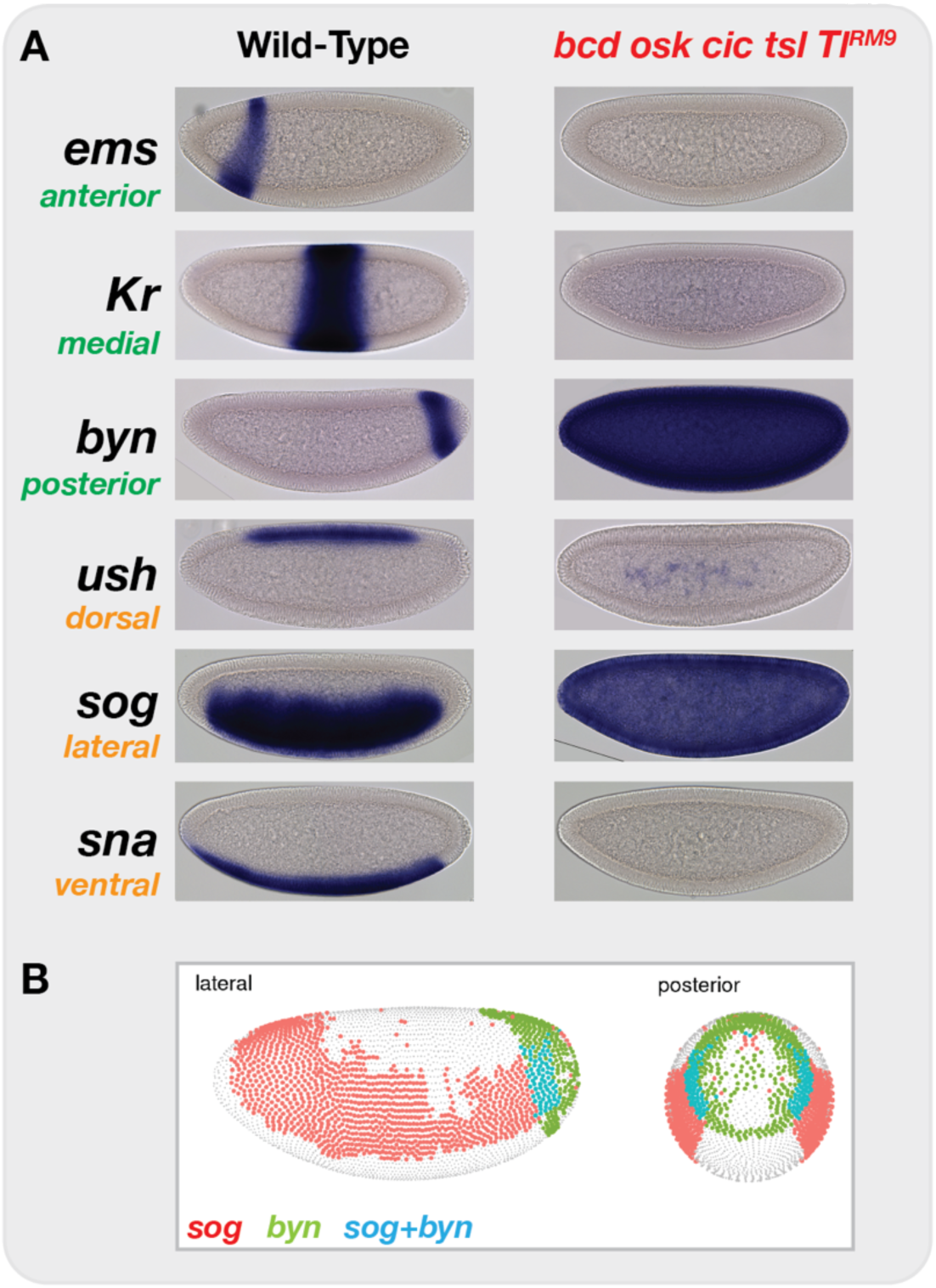
Elimination of graded maternal cues drives development along single uniform lineages. A) in situ hybridization for markers of anterior-posterior (*ems*, *Kr*, *byn*) and dorsal-ventral (*ush*, *sog*, *sna*) marker genes in both wild-type and bcd osk cic tsl TlRM9 mutant embryos. Elimination of graded positional information converts all cells in the blastoderm embryo to posterior-lateral cell types (*sog*+ *byn*+). B) Image from the DVEX virtual expression explorer showing the subset of cells co-expressing both sog and byn.

We therefore performed ATAC-seq comparing single wild-type or mutant embryos precisely staged at 12 and 72 minutes into NC14. As expected, comparison of differentially accessible regions between stage and genotype identifies regulatory elements with spatially restricted expression patterns. For example, at the *wingless* (*wg*) locus, two closely apposed regions (*wg -2.5* and *-1*) show differential behavior between genotypes. The *wg -2.5* region undergoes a modest increase in accessibility in wild-type embryos but not in mutants (Figure 3A, green shading). In contrast, the neighboring *wg -1* region shows greatly increased accessibility in mutant samples (Figure 3A, cyan shading). These two regions both comprise a single previously identified CRM (*wg WLZ4L*) controlling early *wg* expression in both a segment-polarity multi-stripe pattern as well as a single posterior endodermal stripe [46]. On the basis of our ATAC data, we individually cloned the *wg -2.5* and *-1* regions into reporter constructs and compared expression between wild-type and mutant embryos. *wg -2.5* becomes active just prior to gastrulation in wild-type but not in equivalently staged mutant embryos, and exclusively drives expression of the segment-polarity multi-stripe pattern (Figure 3B and data not shown). In constrast, *wg-1* is active earlier in NC14 and exclusively drives expression of the endodermal stripe pattern. As expected, whereas *wg -1* expression is restricted to a single posterior endodermal domain in wild-type embryos, all cells in a mutant embryo have strong *wg -1* expression (Figure 3B). We therefore performed differential enrichment analysis using DESeq2 [47] to identify the complete set of regions with patterning-dependent or -independent changes in chromatin accessibility.

**Figure 3:**
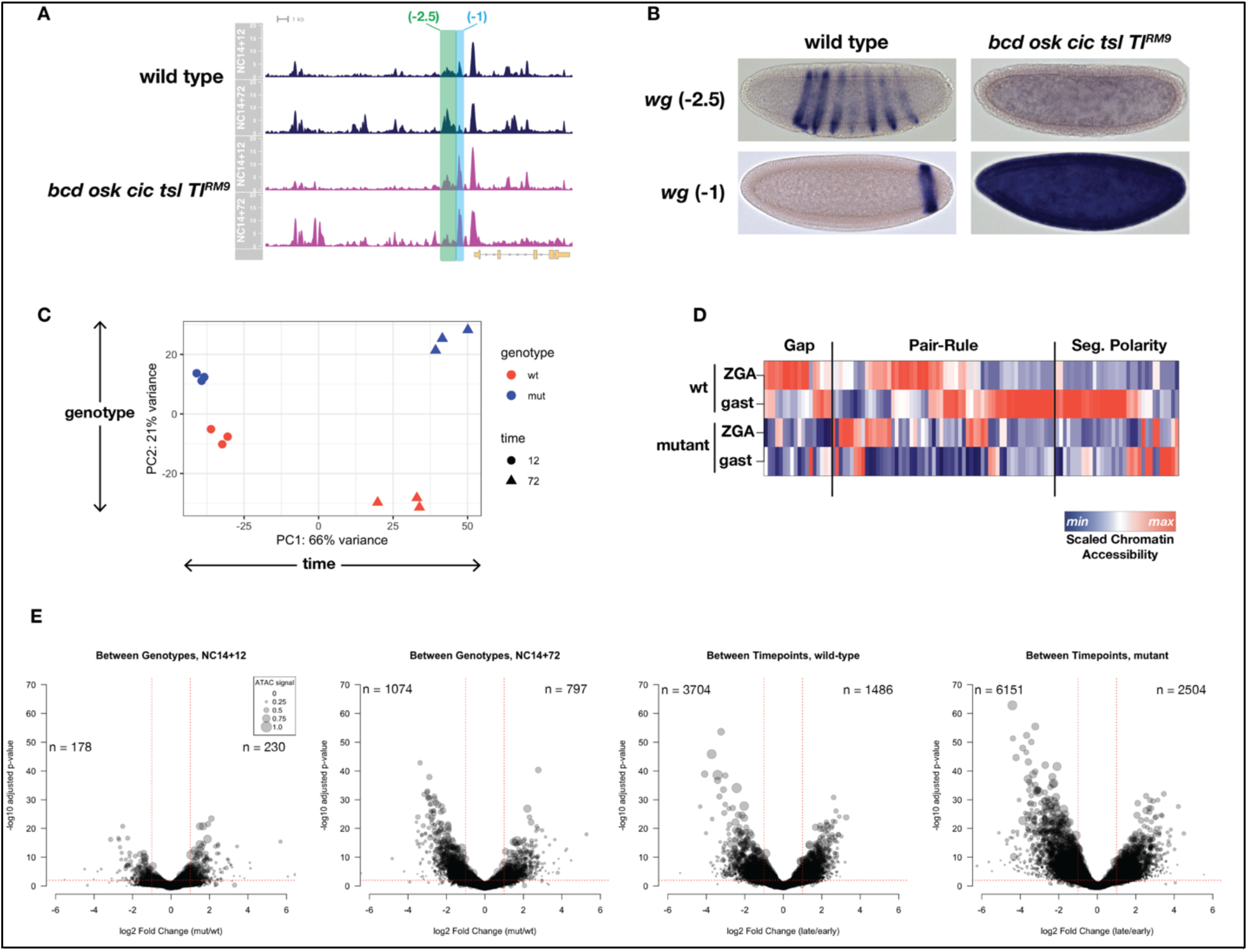
ATAC-seq on wild-type and uniform-lineage embryos resolves time and patterning-dependent changes in chromatin accessibility. A) ATAC-seq coverage over the extended wingless locus. Highlighted regions show closely apposed CRMs with differential responses to patterning inputs. B) Reporters for the CRMs highlighted in A demonstrate separable regulatory inputs into the wg locus, and differential regulation in uniform-lineage embryos. C) Principal component analysis of dynamically accessible regions demonstrates the relative contribution of uniform (time) and patterning (genotype). D) Heatmap showing the scaled chromatin accessibility between ZGA and gastrulation for wild-type and mutant embryos over the complete set of known enhancers within the gap, pair-rule, and segment polarity gene regulatory networks. Colorbar indicates the scaled degree of chromatin accessibility plotted in each column. E) Volcano plots showing effect sizes for comparisons (from left to right) between genotypes early and late, and between timepoints in wild-type and mutant samples. The number of peak regions above significance (0.01) and log2 fold-enrichment (1) thresholds is indicated.

We identify significant sources of both pattern-independent as well as patterning-dependent changes in chromatin accessibility over the period between ZGA and gastrulation. In general, a greater fraction of the sites with dynamic chromatin accessibility undergo patterning-independent changes. This was quantified this in two ways. First, we performed principal component analysis (PCA) on the complete set of differentially accessible regions (i.e., all regions with both a ± 2-fold change and an adjusted p-value > 0.01 either between timepoints or between genotypes as determined by DESeq2). The greatest source of variance is patterning-independent, with the first principal component separating samples according to developmental time (PC1: 66% variance, Figure 3C). The second principal component separates samples according to genotype and therefore resolves patterning-dependent variance (PC2: 21% variance, Figure 3C). There is less of a patterning-dependent difference between NC14+12’ samples compared with the +72’ timepoint, supporting the conclusion that cells initiate the zygotic phase of development with a large degree of chromatin state homogeneity and that heterogeneity emerges over the period leading up to gastrulation from both patterning-dependent and -independent sources.

To relate the observed changes to a single, discrete developmental process, we returned to the set of known segmentation network CRMs and plotted scaled chromatin accessibility over time between wild-type and mutant samples. Within all three tiers of the segmentation network, we find evidence for extensive patterning-dependent chromatin accessibility, at both early and late timepoints (Figure 3D). As previously shown, the gap gene network receives extensive patterning-dependent chromatin accessibility cues from a pioneer activity of Bcd (7/18 CRMs, 38.9%) [32]. Early pair-rule CRMs receive both patterning dependent and independent inputs, however the majority of late pair-rule CRMs gain accessibility in a patterning-dependent manner (25/30 pair-rule CRMs with late accessibility, 83.3%). Segment polarity CRMs (e.g., *wg -2.5* and *wg -1*, Figure 3A) likewise have extensive patterning-dependent accessibility states (16/33, 48.5%). Therefore, these results indicate that although overall changes in accessibility tend to occur independently of embryonic patterning, the networks dedicated specifically to embryonic patterning display a disproportionate reliance on patterning systems for determination of their chromatin accessibility states.

Next, we quantified the types of changes in chromatin accessibility that we observed in our analysis. Similar to the PCA analysis, we find fewer patterning-dependent differences at ZGA than at gastrulation (408 versus 1871, Figure 3E, left two panels). In contrast, a greater number of time-dependent differences are observed for both genotypes (5190 for wild-type, 8655 for mutant, Figure 3E right two panels). We note that these numbers represent above-threshold statistical significance for tests on only one of two critical parameters in this experiment, either time or genotype.

To comprehensively classify the types of changes that any single region undergoes over time and relative to patterning inputs, we took an clustering-based approach to identify groups of similarly-behaved regions and then used the output from paired DESeq2 tests to assign regions to each identified category (see Materials and Methods). By this approach, we identify a total of 2917 dynamic regions that classify into one of ten distinct dynamic categories with respect to time and patterning-dependence (Figure 4). Overall, roughly similar numbers of sites gain and lose accessibility over time with 1351 of sites that are open early losing accessibility over time (46.3%, “late repression” classes, Figure 4) and 1446 sites gaining accessibility by the onset of gastrulation (49.6%, “late accessibility” classes, Figure 4). In total we identify 1635 (56.1%) strictly pattern-independent regions that either gain (n = 725 (24.9%)) or lose (n = 910 (31.2%)) accessibility uniformly between ZGA and gastrulation (Figure 4, see *slam* detail and Figure 4 Supplement 1 & 2). The remaining 1282 regions (43.9%) receive inputs from patterning systems. A set of 307 (10.5%) additional regions gain accessibility uniformly by ZGA but undergo patterning-dependent losses in accessibility by gastrulation (Figure 4, and Figure 4 Supplement 3 and 4). In this case, we cannot distinguish whether the patterning-dependent behavior is repressive or instead represents spatially restricted maintenance of accessibility against a uniform repressor. The remaining classes all reflect pattern-dependent behaviors. A total of 254 regions (8.7%) show pattern-dependent early accessibility. This class is split either into regions that lose accessibility by gastrulation (n = 134, Figure 4, see *gt-10* detail and Figure 4 Supplement 5 and 6) or show constitutive patterning-dependent accessibility through gastrulation (n = 120, Figure 4 and Figure 4 Supplement 7 and 8). Finally, 721 regions (24.7%) demonstrate late patterning-dependent gains in accessibility (Figure 4, see *slp1 “5”* detail and Figure 4 Supplement 9 and 10). Patterning-dependent regulation of chromatin accessibility therefore increases by a factor of 4-fold over the course of NC14. While patterning-dependent accessibility at ZGA is limited to 254 regions, over the course of NC14, patterning systems have a significant impact on the embryonic chromatin landscape driving late gains in accessibility at 721 sites and late reductions in accessibility (or maintenance, see above) at an additional 307 sites for a total of 1028 late patterning-dependent regions.

**Figure 4:**
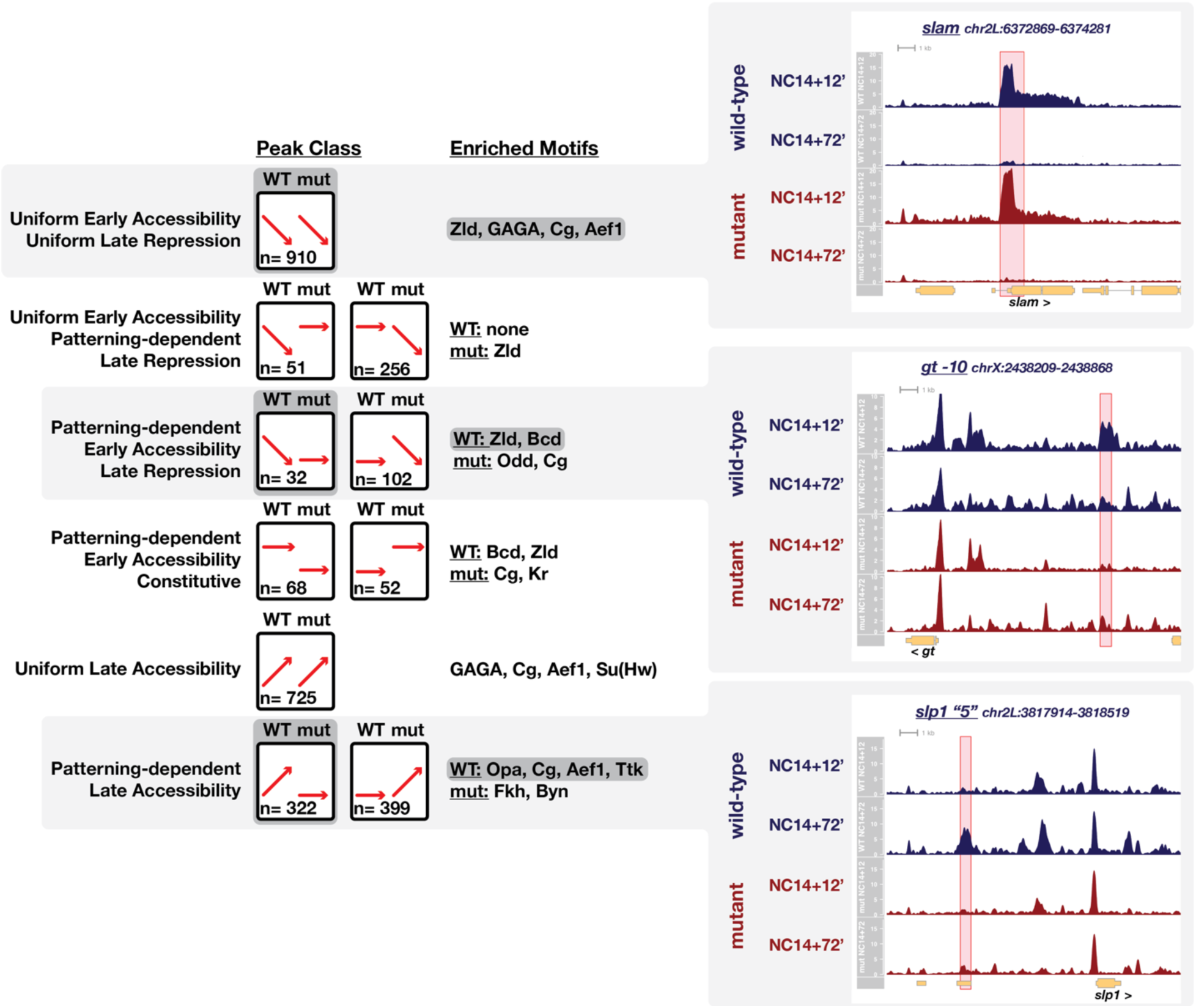
Ten classes of peaks with dynamic chromatin accessibility between ZGA and gastrulation. Ten peak classes are symbolized by cartoons in the left column. Red arrows signify the general behavior of peaks within a class between ZGA and gastrulation for wild-type and uniform-lineage mutant embryos. The number of regions per class is indicated in each cartoon. The center column lists enriched motifs as determined by MEME analysis against a set of 1×104 non-dynamic control regions. The right column shows example ATAC-seq coverage plots for three highlighted classes. The specific peak region within the class is highlighted in red, and we note that additional non-highlighted dynamic regions may be present. Examples of all classes and recovered motifs are provided in the supplemental data.

We find that regions with patterning-dependent regulation are enriched for DNA sequence motifs associated with patterning factors. To gain insight about what regulatory factors could be driving the observed dynamic chromatin accessibility behaviors, we performed sequence motif enrichment analysis within each dynamic class using the MEME suite [48]. Regions with strictly uniform regulation of accessibility are enriched for motifs associated with ubiquitously expressed maternally supplied regulators, including binding sites for Zld [25] and GAGA/CLAMP [41, 49, 50]. In contrast, patterning-dependent sites are enriched for regulators with patterned, spatially restricted expression such as Bcd, Odd-paired (Opa), and Forkhead (Fkh, Figure 4). In addition to these, we frequently observe across all categories enrichment for three maternally supplied factors with expected repressive activity, Tramtrack (Ttk), Adult Enhancer Factor 1 (Aef1), and Combgap (Cg). While Ttk has long been hypothesized to play a broad repressive role over the maternal-to-zygotic transition [35] as well as in regulation of embryonic patterning [51–54], much less is known about potential early embryonic roles of Aef1 and Cg (see Discussion). We also note that although motifs for Cg are not included in the available DNA binding motif databases used to compile these results, we include Cg here on the basis of previous identification of Cg binding to a (CA)_n_ motif [55], maternal expression of Cg, and frequent recovery of an orphan motif (CA)_n_ in our analysis.

The recovery of Bcd and Fkh motifs in our dataset suggests that enrichment analysis could identify potential patterning-dependent pioneer activities responsible for driving differential accessibility states. We note here that recovery of motifs in our analysis is similar to those recovered in another recent report [56], which measured changes in accessibility between early and late NC14 samples but did not distinguish between patterning-dependent and independent events. Bcd has been demonstrated to pioneer accessibility at a subset of its targets [32], and Bcd-motif enrichment in this analysis correlates with the set of previously identified bcd-dependent regions (e.g., *gt -10*). Fkh is the *Drosophila* homolog of a well-characterized pioneer factor FoxA1/2 that operates in early mammalian endodermal development [57–59]. Fkh is expressed zygotically late in NC14 within the posterior endodermal precursors we enrich with mutant embryos, and Fkh motif enrichment is observed specifically within the set of regions that have enhanced late accessibility in mutant embryos (Figure 4). Therefore, it is possible that like its mammalian counterpart, Fkh may pioneer accessibility of distinct endodermal CRMs in early *Drosophila* development.

Within the set of regions with patterning-dependent late accessibility, we also enrich for a zygotic pair-rule transcription factor, Opa (Figure 4). Opa is a C2H2 zinc-finger transcription factor that is the Drosophila homolog of Zn-Finger of the Cerebellum (Zic) proteins, which have been implicated in a broad range of developmental functions ranging from maintenance of stem cell pluripotency, neural crest specification, and neural development [60–64]. Broad roles for Opa in *Drosophila* development are also likely. In addition to an early requirement of Opa for segmental patterning, Opa is necessary for midgut morphogenesis [65], imaginal disc and adult head development [66], and has recently been shown to play a critical role in regulating the temporal identity of neural progenitors [67]. Although neither Opa nor its homologs have not been shown previously to pioneer chromatin accessibility, mouse Zic2 has been demonstrated to bind both active and poised enhancers in embryonic stem cells prior to Oct4 and to play a critical role in maintenance of stem cell pluripotency [61], and in certain species of moth flies Opa has been adopted as the maternal anterior determinant [68], which in *Drosophila* requires pioneer activity from Bcd [32]. This, coupled with the observed enrichment of Opa motifs in our analysis, raised the possibility that Opa could contribute to patterning-dependent chromatin regulation after ZGA.

### Opa pioneers chromatin accessibility of late patterning-dependent regions

Originally identified in a genetic screen for regulators of embryonic segmentation [69], *opa* functions as a pair-rule gene to pattern alternating segmental identities. However, *opa* differs from the other seven pair-rule genes in several ways. First, *opa* is not expressed in a characteristic early seven-stripe pattern typical of pair-rule genes but is instead expressed uniformly over the entire embryonic segmental primordium [70](Figure 5A). Second, compared with other pair-rule genes, *opa* initiates expression significantly later in development and has been proposed to function as a temporal switch within the pair-rule network, facilitating the transition from early to late phases of network operation [71]. During this transition, new regulatory interactions between network components are observed (e.g., emergence of repressive interactions at gastrulation between *paired* and *odd-skipped* within cells where both factors are co-expressed at earlier timepoints) [71], and the mechanism whereby Opa mediates these new network interactions is unclear. Because we observed enrichment of Opa binding motifs in the set of regions that gain patterning-dependent accessibility late in NC14, and because many pair-rule and segment polarity CRMs also display late patterning-dependent accessibility, we tested the hypothesis that Opa functions as a pioneer factor to confer changes in chromatin structure downstream of maternal patterning cues.

**Figure 5:**
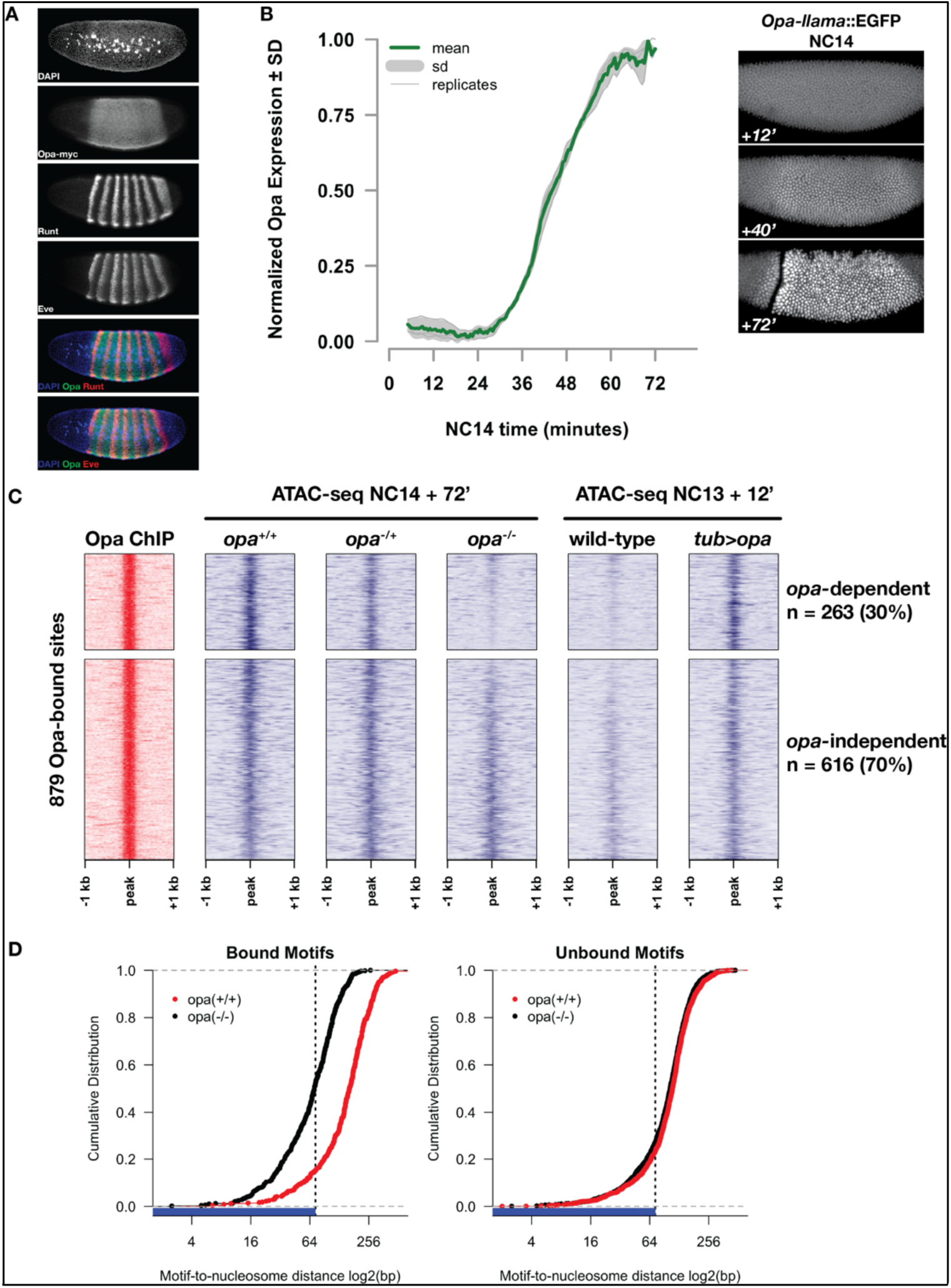
Opa is necessary and sufficient to pioneer accessible chromatin. A) Immunostaining for a myc-tagged opa allele relative to expression of pair-rule genes runt and eve in a late NC14 embryo. B) Opa expression dynamics were measured using a llama-tagged opa allele. Opa expression initiates midway through NC14 and reaches steady levels by entry into gastrulation at 65 minutes. Images show representative expression of opa-llama::EGFP at the indicated timepoints. C) Heatmaps showing scaled ATAC-seq accessibility measurements (blue) over a set of high-confidence Opa binding sites, as determined by ChIP-seq for opa-myc (red). Two experiments are shown, for loss of function at NC14 + 72’, and gain of function at NC13 +12. Loss of blue signal indicates a reduction in accessibility. D) Cumulative distribution of measured of distances between bound (left) or unbound (right) Opa motifs and modeled nucleosome dyad positions, in the absence (black) or presence (red) of Opa. X-axis is log2 scaled, and the expected coverage of a nucleosome is depicted by the blue rug and vertical dotted line.

By ChIP-seq for RNA Pol2 [72], whereas typical pair-rule genes (e.g., *ftz*) show robust Pol2 association as early as NC12, little if any Pol2 is detected at the *opa* locus until NC13, when a small peak of Pol2 forms at the promoter. Productive elongation of *opa* becomes apparent at the beginning of NC14 (Figure 5 Supplement 1). To measure *opa* expression dynamics, we used CRISPR to insert an anti-GFP llama tag [73, 74] to the 3’ end of the *opa* coding sequence and live-imaged *opa-llama* expression in embryos co-expressing free maternal EGFP. Engineered *opa*-*llama* flies are homozygous viable and show no detectable adverse effects from manipulation of the *opa* locus. Consistent with RNA Pol2 measurements, no Opa protein is detected prior to NC14 (data not shown). Llama-tagged Opa first becomes detectable above background by live imaging at 35 minutes into NC14 reaching an apparent steady-state expression level at 60 minutes, shortly before gastrulation (Figure 5B). These measurements indicate that Opa expression is consistent with an exclusively late NC14 role in regulating gene expression over nearly all cells of the embryonic segmental primordium.

By measuring chromatin accessibility in single *opa^+/+^, opa^-/+^,* and *opa^-/-^* embryos at NC14 + 72’, we find that Opa is necessary to pioneer chromatin accessibility at a subset of its direct genomic targets (Figure 5A). We first determined a set of high-confidence direct Opa binding sites by performing ChIP-seq on an engineered allele of *opa* in which we introduced by CRISPR a *3x-myc* epitope tag into the 3’ end of the *opa* coding region. The resultant *opa-myc* allele is homozygous viable, is expressed within the expected domain (Figure 5A) with expected kinetics, and has no detectable adverse effects from engineering of the *opa* locus (data not shown). We performed ChIP-seq on three independent biological replicates of 200 homozygous *opa-myc* cellular blastoderm embryos using as a negative control wild-type (*w^1118^*) embryos. Mapped reads were subjected to peak calling with MACS2 [75] and we defined a set of 879 reproducible, high-confidence Opa ChIP peaks by Irreproducible Discovery Rate analysis [76] (see Materials and Methods). To test whether Opa pioneers accessibility at its direct targets, we performed ATAC-seq on single NC14 + 72’ embryos collected from a cross between parents heterozygous for a null allele (*w; opa^IIP32^/His2Av-GFP*). Following mapping of reads, zygotic genotypes were called based on recovered SNPs (see Materials and Methods). DESeq2 analysis of differentially accessible regions between genotypes determines that 263 (29.9%) of the high-confidence Opa binding sites show significantly reduced chromatin accessibility in homozygous *opa*-mutant embryos (Figure 5C). Therefore, Opa is necessary for conferring accessible chromatin states at a subset of its direct binding sites. Relative to the set of patterning-dependent and independent dynamic genomic regions defined above, opa-dependent pioneer activity alone is sufficient to account for 36% of the set of chromatin regions that depend on inputs from maternal patterning systems to gain accessibility by the onset of gastrulation (Figure 5 Supplement 2). Therefore, in addition to pioneering open chromatin in late NC14 embryos, Opa’s zygotic pioneer activity accounts for a significant proportion of overall changes in chromatin accessibility downstream of maternal patterning systems.

In addition, Opa is sufficient to pioneer open chromatin at the majority of these direct, opa-dependent targets. Because *opa* is expressed late in NC14, at earlier timepoints opa-dependent regions represent effectively naïve chromatin states that can be used to test for sufficiency. We reasoned that at NC13 (one cell cycle before NC14) chromatin accessibility status at Opa targets would largely resemble late *opa-*mutant states. To test for sufficiency, we performed ATAC-seq comparing single wild-type embryos collected 12 minutes into NC13 with embryos misexpressing *opa* maternally under direct control of the maternal *alpha-tubulin 67c* promoter (*tub>opa*) (Figure 5C). Whereas wild-type NC13 + 12’ accessibility at direct Opa targets is nearly indistinguishable from NC14 + 72’ *opa*-mutant samples, maternal expression of Opa increases the open chromatin signature at these targets at 74.9% (197/263) of opa-dependent sites and at 59% (519/879) of direct Opa targets overall. We conclude that, with the exception of 25.1% (66/263) opa-dependent but maternal-opa-insensitive sites that Opa is sufficient to pioneer open chromatin at its target sites.

A second criterion for classifying a transcription factor as a pioneer is that it binds to its DNA motifs even in the nucleosome-associated state [77]. To test this, we measured the distribution of Opa binding motifs relative to nucleosome positions modeled from the distribution of large (>100 bp) ATAC-seq fragments [78]. Either before expression of *opa* (wild-type NC14+12’) or in *opa* mutants at late NC14 (*opa* NC14 + 72’), 51.4% (250/486) of Opa motifs in opa-dependent regions are located within 73 bp of a modeled nucleosome dyad (i.e., within the wrap of DNA around a nucleosome, Figure 5D and data not shown). The fraction of Opa motifs within opa-dependent regions overlapping with predicted nucleosome positions is significantly greater than is observed from the set of Opa motifs located in non-bound, but accessible regions (27.1%, 427/1572, Figure 5D). Following binding of Opa (wild-type NC14+72’), the fraction of motifs in bound regions overlapping the nucleosome footprint reduces to 15% (73/486) and the average position of Opa motifs relative to predicted nucleosome dyads increases by an average of 79 bp in bound regions (Figure 5D). In constrast, the distance between motifs and nucleosomes distance changes by only 6 bp in non-bound regions. These results suggest that a substantial fraction of Opa motifs within future Opa-occupied sites are associated with nucleosomes in the ZGA chromatin state. Following activation of Opa late in NC14, binding of Opa to targets results in a reorganization of nucleosomes surrounding Opa motifs and exposure of previously nucleosome-occluded DNA. Although additional studies of Opa binding to nucleosome-free and -bound DNA will need to be performed, these results are consistent with a model where Opa can interact with nucleosome-associated binding motifs and can trigger reorganization of local chromatin structure.

### Additional mechanisms regulate developmental competence of late segment-polarity CRMs to gain accessibility

We have demonstrated that following ZGA, both local and global changes in chromatin accessibility patterns continue to take place. The identification of pioneers like Opa raises the possibility that distinct zygotic factors function to establish accessibility states conditional on prior, maternal patterning information. We wished to investigate the hypothesis that the sequence of chromatin accessibility changes are themselves critical for the proper execution of the developmental patterning program. To address this, we therefore quantified opa-dependent chromatin accessibility within the segmentation network and evaluated the consequences of premature *opa* expression on patterning. We predicted that maternal mis-expression of *opa* would effectively conflate a ZGA chromatin state with a gastrulation chromatin state. Opa is necessary for full chromatin accessibility at a set of late pair-rule and segment polarity CRMs (Figure 6A-C). Late-acting opa-dependent CRMs within the pair-rule network include the late *eve* seven-stripe element (*eve-late* also referred to as *eve-autoregulatory*,[79–81], see also Figure 6 Supplement 1), the ‘center cell’ repressor that splits early *prd* stripes into anterior and posterior stripes (*prd cc repressor* [82]), a late anterior stripe repressor for *prd* (*prd A8 repressor* [82]), several late *slp1* enhancers (*u3525, i1523, u4739,* and *‘5’*, [83, 84] see also Figure 6 Supplement 1), as well as a novel *odd* late enhancer (*odd-late*, Figure 6B, D and E). Several segment polarity CRMs are also opa-dependent, including a *gsb* 3’ enhancer ([85], Figure 6C), a portion of the *en H* regulatory element (*en H2* [86], see Figure 1), an *en* intronic enhancer [86] as well as less well characterized elements within *ptc* and *wg* defined by large-scale enhancer screens (*ptc 5’* = GMR69F07, *ptc 5’* (2) = GMR69F06, *wg intron* = GMR16H05)[87]. We confirmed an effect of *opa* loss of function on the expression patterns of three opa-dependent CRMs, *odd-late*, *eve-late*, and *slp1 “5”* (Figure 6B, D and Figure 6 Supplement 1). In *opa* mutants, no expression is seen from *odd-late* (Figure 6D). The effect of *opa* on *eve-late* is also nearly complete, although reduced levels of expression persist within stripe 1 (Figure 6 Supplement 1). Loss of *opa* also reduces expression from *slp1 “5”*, completely eliminating activity within secondary odd parasegmental stripes, and significantly reducing activity within the primary even parasegmental stripes (Figure 6 Supplement 1). These effects are largely consistent with the range of previously reported *opa* loss of function phenotype on pair-rule and segment polarity targets [71] and supports the conclusion that the primary function of *opa* in the segmentation network is to modulate the temporally restricted accessibility of a subset of critical CRMs.

**Figure 6:**
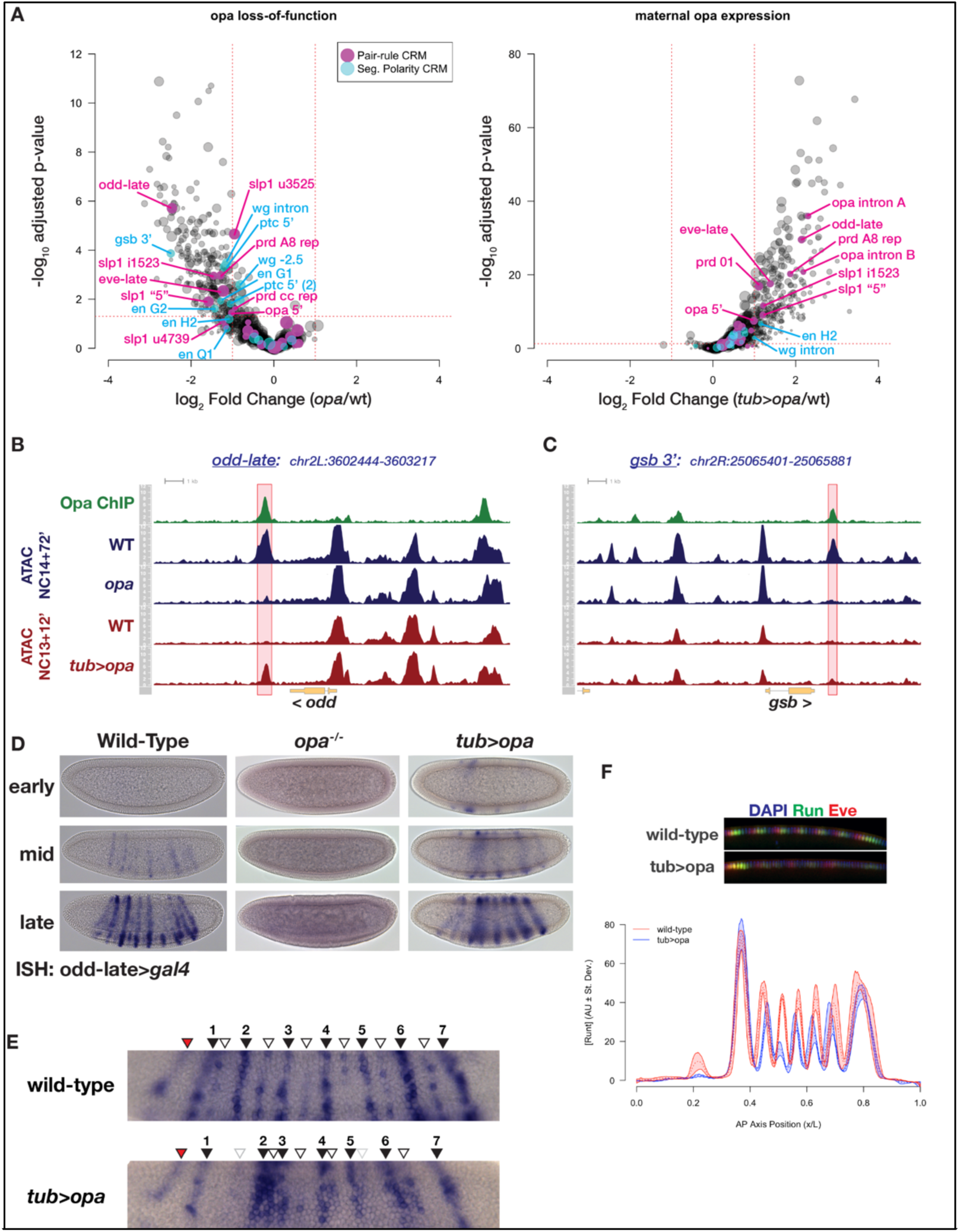
Premature expression of Opa disrupts pair-rule patterning. A) Volcano plots for loss of function (left) and maternal misexpression (right) of Opa with pair-rule and segment polarity CRMs highlighted (magenta and cyan, respectively). B) Example of ATAC seq coverage over an opa-dependent pair-rule locus, odd-skipped. The odd-late CRM is highlighted showing the effect of opa at this element. C) For comparison, ATAC-seq coverage over a segment polarity locus, gooseberry, is shown. The gsb 3’ CRM is highlighted. D) In situ hybridization for an odd-late gal4 reporter is shown for wild-type, opa mutant and tub>opa. Embryo stages are indicated at left. Note the lack of activity in opa mutants and the premature activation in the presence of tub>opa. E) Detail view of odd-late expression in wild-type and tub>opa gastrula stage embryos. Odd parasegmental stripes are indicated with numbered black arrowheads, even stripes with open arrowheads, and the anterior head stripe is indicated with a red arrowhead. Weak even parasegmental stripes in tub>opa that eventually appear are indicated with grey open arrowheads. The stripe at numbered position 1 is coincident with the cephalic furrow, which is beginning to form in both pictured embryos. F) Maternal opa interferes with stripe positioning and intensity for pair-rule genes eve and runt. Plot shows the average effect of maternal opa on runt expression in mid NC14 embryos. Average quantified Runt expression ± std. dev. is plotted for wild type (red) and tub>opa (blue, n= three embryos per genotype). Inset shows dorsal mid-saggital view of a representative embryo of the indicated genotypes stained for Runt (green) and Eve (red).

Notably, although *opa* is sufficient to induce accessible chromatin at most of the late pair-rule opa-dependent targets (*odd-late, eve-late, slp1 “5”, slp1 i1523, prd 01, prd A8 repressor*) as well as three regions within the *opa* locus itself, segment polarity targets show a distinctly reduced sensitivity to gain accessibility in response to premature *opa* expression, with only *en H2* and an intronic *wg* region having marginally above-threshold increased accessibility. To characterize this Soluri et al.: Patterning-dependent chromatin accessibility further, we performed motif enrichment analysis over the entire set of opa-dependent, opa-insensitive sites and find enrichment of the Opa motif, as well as enrichment of two maternal repressors, Ttk and Cg. In contrast, the set of both opa-dependent and opa-sensitive regions shows enrichment for the Opa motif alone. These results indicate that within the segmentation network there exists differential sensitivity to acquisition of chromatin accessibility states, and that although Opa is necessary for conferring open chromatin at a set of pair-rule and segment polarity loci, it is possible that competence to respond to Opa, and perhaps any pioneer in general, is further regulated by additional repressive mechanisms.

Maternal expression of *opa* triggers premature activity of opa-dependent targets and mispatterning of the pair-rule network (Figure 6D-F). The opa-sensitive target *odd-late* initiates expression in wild-type embryos in a domain of seven odd-numbered parasegments that correspond to the secondary pattern of *odd* expression (Figure 6D, E) [71]. Expression within even-numbered parasegments corresponding to the primary *odd* expression pattern follows later, ultimately yielding a 14-stripe pattern (Figure 6E and data not shown). In addition to the parasegmental expression pattern, there is a transient head stripe anterior to the first stripe (Figure 6E). Activity of this CRM is entirely dependent on *opa* (Figure 6D). Maternal expression of *opa* affects both spatial and temporal aspects of the *odd-late* expression pattern, driving premature activity of *odd-late* in stripes with incorrect spatial distributions. The most significant spatial effect of maternal *opa* expression on *odd-late* pattern is an increased interstripe distance between the first and second odd parasegmental expression domain, resulting in apparent compression between the second and third stripes (Figure 6E). In addition, compared with wild-type, the regular spacing of odd- and even-parasegmental stripes at positions 4-6 is disrupted with maternal *opa* expression. Despite the fact that *tub>opa* is expressed uniformly across the entire embryo, premature *odd-late* expression appears not uniformly, but within the spatial domains to which its later expression will be restricted (Figure 6D). Similar effects have been observed with premature expression of *opa* using the *gal4*-UAS system, where increasing levels of *opa* expression lead to defects in *slp1* expression only within the segmental primordium (i.e., in a domain where additional co-regulators are expressed). Only in combination with uniformly misexpresssed *runt+opa* are ectopic domains of *slp1* observed outside the segmental primordium [88]. These observations are consistent with a role for *opa* largely restricted to regulating accessibility and not strictly activating or repressing expression from direct targets such as *odd-late* and that interfering with timing of CRM accessibility results in misinterpretation of regulatory cues and disruption of the precision of embryonic patterning.

To quantify the effect of maternal *opa* expression on additional components of the pair-rule network, we immunostained embryos for Runt and Eve and quantified stripe positioning in precisely staged embryos [89]. Although between genotypes the segmental primordium is unchanged in both overall area and overall positioning, the intensity and positions of stripes 2-6 are affected by maternal *opa* expression. Similar to *odd-late* we also observe increased inter-stripe distances between Runt stripes 1 and 2, and reduced distance between stripes 2 and 3, as well as minor mis-positioning of stripes 4-6. Despite these effects on stripe positioning and intensity, embryos produced from mothers heterozygous for *tub>opa* hatch and can mature to adulthood, albeit with varying degrees of segmental mis-patterning. We note that the long-term severity of the *tub*>*opa* phenotype may be attenuated by limited effects of maternal *opa* expression on the segment polarity network. The positioning of stripes in the pair-rule network is highly precise and results from optimal decoding of maternal patterning inputs [16]. Our results are consistent with a model where premature *opa* expression alters how maternal information is interpreted at the level of this network. While we cannot at present deconvolute effects on the system that stem from Opa-dependent pioneering from possible Opa-dependent effects on transcription, these results provide support for the hypothesis that the sequential transitions in chromatin accessibility patterns are themselves necessary for proper patterning of the embryo.

## Discussion

To the extent that the developmental program can be described by gene regulatory networks, mechanisms for the regulation of chromatin accessibility must play a major role in determining how this program unfolds. We have demonstrated that post-ZGA changes in chromatin accessibility patterns are encoded in the developmental program at multiple levels and have provided evidence that the sequence in which changes in accessibility patterns are introduced to the embryonic chromatin landscape is itself important for the proper interpretation of developmental cues.

One of the most intensely studied developmental transitions in chromatin structure is large-scale ZGA that accompanies the maternal-to-zygotic transition. Our observation of continued uniform, patterning-independent changes to chromatin accessibility after ZGA suggests that additional global timers of developmental progression continue to operate following the maternal-to-zygotic transition. The initial ground state of chromatin structure is built by maternally supplied pioneers, particularly Zelda [7, 25–28, 30], and likely a combination of factors that bind (GA)_n_ repeats, GAGA-factor and CLAMP [7, 41, 72, 90]. Prior studies on the temporal dynamics of the establishment of the initial ground state have suggested that these pioneers act in distinct temporal waves, with the earliest accessible regions associated with Zelda binding, and the latest accessible regions enriched for the (GA)_n_ motif and GAGA-factor binding [7]. Our results add to these prior observations by suggesting that these global ‘waves’ of accessibility regulation continue well past the ZGA, and likely, based on motif enrichment, receive inputs from additional maternally-supplied factors with expected repressive activity: Ttk, Aef1, and Cg. Next to nothing is known about the role of these factors specifically in the context of global chromatin accessibility regulation at the ZGA. Aef1 was identified as a Zn-finger transcriptional repressor that regulates gene expression in adult *Drosophila* fat body [91]. Cg has been implicated in both positive and negative regulation of target genes, has been shown to interact with GAGA factor [92], and plays a role in recruitment of polycomb group proteins to polycomb response elements through direct binding of (CA)_n_ repreats [55]. Both Aef1 and Cg are expressed maternally, and transcripts are cleared by the onset of gastrulation (Berkeley Drosophila Genome Project) although protein expression kinetics have not been determined for early embryos. The transcriptional repressor Ttk plays a role in the spatio-temporal regulation of segmentation gene expression, directly influencing timing and pattern of pair-rule and segment polarity gene expression [35, 51–54]. Perhaps indicating a more general role beyond segmentation network regulation, Ttk has been demonstrated to interact directly with GAGA factor and antagonize GAGA-dependent transcriptional activation [92, 93]. Because Ttk mRNA and protein are expressed at high levels maternally and are cleared from the embryo by gastrulation [51], Ttk (as well as Aef1 and Cg) may play a more global role in ZGA timing by limiting pioneer factor activity at target sites until they are cleared from the embryo. In this respect, it is interesting to note that the subset of opa-dependent targets that are insensitive to maternal *opa* expression demonstrate an enrichment for Ttk and Cg motifs. One possible explanation for this apparent developmental competency to respond to Opa pioneering activity is that binding of maternal repressors can antagonize pioneer factor activity. Future work will include testing the role of these maternal factors in the context of ZGA timing and regulation of coordinated, global chromatin remodeling events.

In contrast with these mechanisms for uniform regulation of accessibility, while there is relatively little influence of maternal patterning systems directly on chromatin accessibility status, we observe that certain zygotic targets of maternal pathways, such as Opa, can have a major impact on chromatin accessibility states. Opa’s primary role in the pair rule network is to facilitate the transition, termed a ‘frequency doubling’, from early to late expression patterns. In the absence of Opa, pair-rule loci (primarily *odd*, *slp, run,* and *prd*) fail to undergo the transition from early seven-stripe to late 14-stripe patterns. Additionally, late 7-stripe expression of *eve* is also strongly affected [71]. Because of uniform expression across the segmental primordium, Opa does not provide positional information that defines the precise location of its target expression domains [70], and has been proposed to cooperate with additional pair-rule factors such as Runt to activate or repress target gene expression [71, 88]. Here, we demonstrate the mechanism for Opa’s role in the network: that Opa facilitates the frequency doubling of the pair-rule network by pioneering accessibility of the CRMs that drive these late expression patterns. We predict that Opa pioneer activity will therefore result in conditional cis-regulatory interactions of the remaining pair-rule factors with late CRMs. This mechanism can help explain the previously observed ‘conditional regulation’ between network components (e.g., Odd repression of *prd* to yield anterior and posterior stripes)[71], which we propose is largely mediated through opa-dependent CRM accessibility states. The set of opa-dependent CRMs within the pair-rule network that we identify strongly support this conclusion. Incorporating such ‘time-gated’ pioneering events into a regulatory network may therefore allow for a system to generate multiple patterning outputs from a limited set of input transcription factors. Further investigation of the opa-dependence for conditional cis-regulatory interactions amongst pair-rule factors, as well as identification of additional zygotic pioneer factors will address these predictions.

A critical distinction that arises between transcription factors within a network, then, is what effect they have on chromatin accessibility states. It is likely that not all transcription factors have pioneer activity, or that the ability of a factor to pioneer is context specific. For instance, loss of *grainyhead* (*grh*) has minimal effects on the pre-gastrula chromatin accessibility state, despite the fact that *grh* has been demonstrated to function as a pioneer in other biological contexts [56]. Similarly, while repressors have been demonstrated to negatively impact chromatin accessibility states, certain repressors, such as the pair-rule factor *hairy*, can operate not through compaction of chromatin but by inhibiting recruitment of the basal transcriptional machinery, at least in certain contexts [94]. Whether repressors within the pair-rule network fall into distinct chromatin-dependent and -independent categories at a genome-wide scale remains to be determined. A comprehensive appraisal of how transcription factors within a network not only interact with cis-regulatory elements over time but also how they impact chromatin accessibility states will be necessary to fully understand the regulatory logic of embryonic patterning.

### Is accessibility regulated maternally along the DV axis?

We note that we have not yet exhaustively examined all possible maternal patterning contributions to chromatin accessibility. The *bcd osk cic tsl Tl^RM9^* mutant embryos used in this study, while amorphic for maternal determinants of AP and terminal cell fates, have uniform, moderate *Tl* pathway activation and therefore moderate *dorsal* (*dl*) activity [44]. So, it remains possible that any possible *dl*-dependent chromatin accessibility states remain unidentified by our study. However, we predict that *dl* does not pioneer chromatin at least to the same extent as *bcd*. The observation of *bcd*-dependent chromatin accessibility states has now been observed in four independent studies in addition to this work [8, 32–34]. Besides studies where accessibility was measured in *bcd* mutants directly (this work and [32]), two additional studies of spatially restricted chromatin accessibility independently confirmed the predicted [32] anterior enrichment of *bcd*-dependent regions [33, 34]. However, none of these studies were specifically designed to distinguish differential DV accessibility states. In contrast, a recent single-cell ATAC-seq study could have identified these states if they existed [8]. This study consistently identified several clusters of early embryonic cells that distinguished AP states, and single-cell ‘anterior’ clusters were found to be enriched for previously identified *bcd*-dependent regions [8]. If *dl* were to pioneer chromatin to a similar extent as *bcd*, we would therefore expect that single-cell ATAC-seq would have also identified early “dorsal” and “ventral” clusters, but in this study, all DV-specific clusters (e.g., mesoderm) were only found associated with cells presumed to be staged later based on ‘pseudotime’ analysis. So, although maternal systems may not drive differential accessibility along the DV axis, it nevertheless likely that, similar to *opa*, specific zygotic targets within the DV networks operate immediately downstream of maternal pathways to distinguish DV-specific chromatin accessibility states. This also highlights how in the *Drosophila* system both single-cell (or enriched cell) methods and genetic manipulation represent powerful complementary approaches for distinguishing the degrees of complexity in the chromatin landscape and linking these features to developmental regulatory systems. Replacing *Tl^RM9^* with complete gain- or loss-of function alleles affecting *Tl* signaling [95] in our maternal mutants will definitively test the hypothesis suggested by single-cell approaches of limited early DV heterogeneity in chromatin accessibility states.

### Transcriptional activation or pioneering?

A prior study [34] observed that chromatin accessibility levels at enhancers as measured by ATAC-seq correlated strongly with the transcriptional activity of the associated gene. This raises important questions about how to interpret ATAC-seq peak intensities, including whether losses of ATAC signal in certain mutant backgrounds can be interpreted as stemming from pioneer activities or simply reflect reduced expression of associated genes. While this observation points to likely several compounding layers of regulation that influence chromatin accessibility and ATAC-seq measurements, it is unlikely that transcriptional activity is the sole determinant of enhancer accessibility states. For one, sites can be found that gain accessibility uniformly, but which are not transcribed. One such locus, *ush*, undergoes significant patterning-independent gains in promoter accessibility between ZGA and gastrulation (Figure 4 Supplement 2), but fails to be expressed in the mutant (Figure 2). This demonstrates that gains in accessibility can occur independently of transcriptional activation, and similarly highlights an example where transcriptional repression need not result in reduced accessibility. In contrast, the high degree of ATAC signal we observe at sites that preferentially gain accessibility in uniform-lineage mutant embryos (e.g. *wg -1* (Figure 3 A, B) and *18w 1* (Figure 4 Supplement 10)) may be significantly compounded by the large fold-increase between wild-type and mutant embryos of cells in which the associated CRM drives active transcription (Figure 3B). So, we cannot rule out completely an additional contribution from transcriptional status on the magnitude of ATAC signals.

In our study, we demonstrate distinct patterning-dependent gains in chromatin accessibility over time, and a significant fraction of these we link to *opa* activity. However, within the segmentation network specific CRMs, Opa is not the sole pair-rule factor predicted to bind to these sites. At the *odd-late* enhancer, for instance, in addition to our demonstration of direct Opa binding, prior ChIP-chip profiling [96] of a subset of pair rule genes in broadly-staged embryos predicts that one activator (Ftz) as well as several repressors (Slp1, Prd, Run, and to a lesser extent Hairy) bind to this region. Ftz, Run, and Hairy are expressed during early NC14, well before Opa, yet neither is *odd-late* functional at this time, nor is it accessible. Our results are consistent with a model where this and other *opa*-dependent enhancers require Opa to pioneer accessible chromatin first in order for other regulatory factors such as Ftz and Run to interact with the underlying DNA sequence and thereby regulate patterned expression of the associated gene. It is possible that robust transcriptional activity following initial pioneering results in additional increases in overall ATAC signal. Future work will include directly testing whether Opa expression facilitates *de novo* interaction of pair-rule genes at previously inaccessible chromatin regions, and direct testing of the informatic consequences of dynamic chromatin accessibility states on network function.

We concede that it is difficult to disentangle using ATAC-seq alone what are likely numerous additional determinants of developmental chromatin accessibility dynamics. For instance, ATAC does not provide good information about the chromatin state of enhancers prior to acquiring an accessible state, nor does it directly shed light on the mechanisms for regulating competence for a site to be pioneered. In the case of Opa, we identify shifts in the positions of nucleosomes relative to binding sites consistent with a model where Opa binding triggers reorganization of chromatin structure at targeted loci. In many, but not all cases, Opa is sufficient to open chromatin at these sites earlier in development. Additional contextual features of the system must regulate competency for a site to be pioneered. As observed previously [33, 34], in many cases regions that undergo spatially restricted, pattern-dependent changes in chromatin accessibility demonstrate intermediate degrees of accessibility in the ‘closed’ state greater than that observed at completely inactive genomic loci. The idea of distinct degrees of ‘repressed’ or ‘closed’ chromatin adds further layers of potential regulation to these systems and raises the possibility that additional factors confer competence for chromatin to be pioneered. Much remains to be understood about what grants competence to a genomic locus to undergo chromatin remodeling in response to binding of a pioneer factor.

## Materials and Methods

### Drosophila Stocks and Husbandry

All fly stocks were maintained on an enriched high-agar cornmeal media formulated by Gordon Gray at Princeton University. Embryos were collected on standard yeasted apple-juice agar plates mounted on small cages containing no more than 300 adults aged no older than 14 days. Media compositions are available upon request.

The wild-type stock for ATAC-seq experiments is *w*; *His2Av-EGFP* (III), and is *w^1118^* for immunostaining and *in situ* hybridizations. The quintuple “patternless” maternal mutant was a gift from Eric Wieschaus at Princeton University. The base stock genotype is *bcd^E2^ osk^166^ cic^1^ tsl^1^ Tl^RM9^* and was constructed through multiple rounds of meiotic recombination. A commonly available *bcd osk* double mutant stock was recombined with *cic tsl* double mutants and resultant quadruple mutants were subsequently recombined with *Tl^RM9^*. Retention of mutant alleles was confirmed at each step by extensive complementation tests against single mutant alleles and examination of cuticle phenotypes. For ATAC-seq experiments, patternless mutant mothers were trans-heterozygotes between two independent quintuple mutant recombinants, and one of the two mutant chromosomes also carried *His2Av-EGFP* to facilitate staging of embryos. The *opa* mutant allele (*opa^IIP32^*) was obtained from the Schupbach-Wieschaus Stock Collection at Princeton University [69]. For ATAC-seq experiments, *opa*/TM3, Sb were first crossed to *w*; *His2Av-EGFP* and embryos were collected from a cross between resultant *opa*/*His2Av-EGFP* adults. In the course of this study, we determined the nature of the *opa^IIP32^* lesion as a single G>A point mutation (chr3R:4853998-4853998, dm6 assembly) that generates a missense allele converting a critical cysteine residue within the first predicted Opa C2H2 Zn-finger to tyrosine (C298Y). Since this lesion is predicted to severely impact Opa DNA binding and mutant *opa^IIP32^* cuticle phenotypes are indistinguishable from those produced from homozygous *opa^Df^* (data not shown), we conclude the *IIP32* allele is effectively amorphic.

### CRISPR modification of the *opa* locus

CRISPR was performed by microinjection of *y sc v*; *nos>Cas9*/*CyO* embryos with ∼60 pl of a mixture of plasmids encoding pU6-driven sgRNAs targeting *opa* and *ebony* (100 ng/ul each) and a plasmid for homology directed repair to insert either a 3x-myc tag or an EGFP-enhancer llama-tag [73, 74] at a concentration of 500 ng/ul. All injections were performed in-house using a modified in-chorion approach originally developed by Nicolas Gompel and Sean Carroll (http://carroll.molbio.wisc.edu/methods/Miscellaneous/injection.pdf) which greatly facilitated high-throughput injection and enhanced survival of hundreds of embryos per day. The cited protocol was enhanced by beveling unclipped, freshly pulled injection needles at a 40° angle on a Narishige Needle Grinder (Narishige, Tokyo, Japan), resulting in reproducible needle preparations that easily pierced the chorion and minimized damage to injected embryos from irregularly surfaced tips. Targeting efficiency of *ebony* was used to select for likely ‘jackpots’ of CRISPR modified founders, as described in [97]. The *opa* sgRNA target sequence was selected from the UCSC Genome Browser/CRISPOR pre-calculated sgRNA sequences [98] and targets chr3R:4869148-4869167 (+ strand, dm6), yielding an expected Cas9-dependent double-strand break 4 bp downstream of the desired insertion site (immediately upstream and in-frame with the *opa* stop codon). The guide sequence oligonucleotides were synthesized with a leading G residue and terminal sequences compatible with insertion into BbsI-digested pU6-2xBbsI as described at http://flycrispr.org/protocols/grna/ [99]. Homology-directed repair (HDR) constructs were constructed using two 500 bp homology arms either terminating at the insertion site on the left or beginning at the expected Cas9 double-strand break on the right. Because the selected *opa* sgRNA target sequence spanned the site of insertion and would therefore be disrupted either in the HDR vector or in successfully targeted and repaired *opa* loci, we did not introduce sgRNA or PAM site mutations to prevent CRISPR targeting of the HDR plasmid or re-targeting of repaired loci. We constructed HDR sequences in a modified pBluescript KS vector where we introduced a *3xP3>dsRed* eye marker into the vector backbone (pBS 3xP3) to facilitate negative selection of unwanted founders that incorporate the full HDR plasmid by homologous recombination. All DNA constructs for injection were prepared using a modified Qiagen EndoFree Midiprep protocol (Qiagen, Hilden, Germany), which enhanced survival of injected embryos compared with standard Qiagen Midiprep grade DNA preparations. Following injection, surviving embryos were raised to adulthood and single founders were crossed to five or six *w; Dr e/TM6c, Sb* flies. F1 progeny were scored for the proportion of *ebony* and vials with >50% *ebony* progeny were scored as “jackpots” in which we induced greater-than-monoallelic targeting of the *ebony* locus. Up to six individual jackpot males from up to six jackpot founders were subsequently crossed to *w; Dr e*/*TM6c, Sb* females and stocks were established from F2 progeny. We favored jackpots that produced 85-100% *ebony* progeny. We observed a 32% survival rate of injected embryos (effectively 64% survival taking into account homozygous lethality of either the second-chromosomal *nos>Cas9* transgene or *CyO*), of these 40-60% of founders produced at least one *ebony* progeny, and 15-25% of surviving founders, overall, produced jackpot lines. Following selection against whole-pasmid integrants (*dsRed+* individuals), approximately 75% of selected F2 jackpot individuals were positive for the desired insertion. No difference in insertion efficiency was observed between *3xMyc* and the *EGFP-llama* cassettes. Positive insertions were identified by PCR and reared to homozygosity before PCR amplification of the modified locus (using primers targeting sites outside the original left and right homology arms) and sequence confirmation by Sanger sequencing. Between two and four stocks representing identical independently generated alleles were kept and confirmed for general stock viability and overall expression patterns/functionality of the modified locus, but core experiments were performed on single representative stocks isogenic for individual engineered chromosomes.

### Time-lapse imaging of *opa-llama* expression

Embryos produced from a cross between *w His2Av-RFP; bcd_promoter_>EGFP/+* females and *w; opa-llama* males are maternally loaded with RFP-marked histones, low levels of free uniformly expressed EGFP, and are heterozygous for a llama-tagged *opa* allele. Embryos were dechorionated for one minute in 4% Clorox (50% dilution of a commercially available “concentrated” Clorox Bleach solution), washed extensively in dH_2_O, and mounted in halocarbon oil sandwiched between a glass coverslip and a gas-permeable membrane mounted on an acrylic slide. Syncytial blastoderm embryos were identified on the basis of morphology and nuclear density, and time-lapse images were collected on a Leica SP8 WLL confocal microscope (Leica Microsystems, Wetzlar, Germany) with a 40x 1.3 NA oil-immersion objective, using excitation wavelengths of 489 nm for EGFP and 585 nm for RFP. Images were captured at 512 x 240 pixels with 692 nm x-y pixel size (0.83x Zoom) and a 1 um z-step spanning a z-depth of 12 um, 115 um pinhole size, and 110 Hz scan rate with bidirectional scanning. Series were collected at 30 seconds per xyz stack. We required that a movie begin within NC12 and for data collection to continue until at least NC14 + 72 minutes. This ensured not only that we quantified *opa* expression from the beginning of NC14, but also controlled for variation in developmental rates by allowing us to measure the duration of NC13. We required that NC13 would last no longer than 22 minutes and no shorter than 17 minutes for imaging to proceed. Fluorescence intensity was quantified by segmenting nuclei in the RFP channel and quantifying average, per nucleus intensity in the GFP channel. Nuclei outside of the *opa* expression domain were manually identified to estimate fluorescence background and plotted expression levels were calculated by subtracting the background estimate from the average intensity of nuclei within the observed *opa* expression domain. Values from three biological replicates (three individual embryos imaged on three different days from two independent crosses) were averaged to yield the plotted data.

### Construction of *tub>opa* for maternal misexpression

*opa* genomic sequences were amplified from a single squashed homozygous adult *opa-3xmyc* fly by standard PCR. Two PCR amplicons containing the entire *opa* coding sequence, omitting the large 14.1 kb first intron but retaining the small 140 bp second intron, were joined in-frame and inserted between the maternal *alpha-tubulin 67c* promoter and *sqh* 3’UTR in the transgenic vector pBABR-tub-MCS-sqh [32]. This yields maternal *alpha-tubulin* driven *opa*-*3xmyc* uniformly across the AP axis. Transgenic lines were generated by injecting 60 pl of a 200 ng/ul solution of the transgenic vector into *y sc v nos>phiC31 int; attP40*, outcrossing founders to *w; Sco/CyO* and selecting for mini-white positive transformants. Stocks were maintained unbalanced because of apparent synthetic dominant maternal effect lethality when maintained in *trans* to CyO that was not evident when stocks were maintained over a wild-type second chromosome.

### ATAC-seq

ATAC-seq was performed essentially as described in [7]. Briefly, embryos were staged at NC14 to the minute by observation of anaphase of the prior cell cycle (t = 0’) and aging embryos to the desired stage. Embryos were maintained at constant room temperature to reduce variability in staging. Three minutes before the desired stage, a single embryo was dechorionated and macerated in Lysis Buffer [100]. Nuclei were pelleted by gentle centrifugation at 4°C at 500 RCF and the supernatant was discarded prior to freezing on dry ice. We have determined that one critical parameter in this process is the use of low-retention 1.5 ml microcentrifuge tubes (e.g., Eppendorf DNA LoBind tubes, manufacturer part number 022431021) (Eppendorf, Hamburg, Germany). Samples were briefly thawed on ice before tagmentation in a 10 ul solution as described previously. A second critical parameter was to ensure sufficient resuspension of the nuclear pellet during the addition of the tagmentation solution, which we achieve by extensive up-and-down pipetting upon addition of the solution. Samples were incubated for 30 minutes at 37°C with 800 RPM agitation in an Eppendorf Thermomixer HotShakey device. Following tagmentation, samples were cleaned up with the Qiagen MinElute Reaction Clean Up kit, eluting in 10 ul.

ATAC libraries were amplified using a set of modified Buenrostro primers that introduced Unique Dual Indexes (UDI). Buenrostro primers differ slightly from Illumina Nextera primers with the addition of 7 bp (Nextera Index Read 2 Primer/ Universal or i5) or 6 bp (Nextera Index Read 1 Primer/i7) additional base pairs to the 3’ ends of standard Nextera primer sequences [100] (Illumina Inc., San Diego, California). Although we have not confirmed this directly, the additional bases in the Buenrostro primer design may yield improved amplification efficiency from ATAC libraries than standard Nextera primers as has been reported for a different Tn5 tagmentation-dependent assay, Cut and Tag [101], see https://www.protocols.io/view/bench-top-cut-amp-tag-z6hf9b6). Therefore, we designed modifications of the original Buenrostro primer design rather than purchase commercially available UDI Nextera primer sets. The generalized UDI ATAC primer sequences are:

ATAC UDI Index Read 2 (i5):

5’-AATGATACGGCGACCACCGAGATCTACACnnnnnnnnTCGTCGGCAGCGTCAGATGT*G-3’

ATAC UDI Index Read 1 (i7):

5’-CAAGCAGAAGACGGCATACGAGATnnnnnnnnGTCTCGTGGGCTCGGAGATG*T-3’

Where the ATAC UDI Index Read 2 primer corresponds to the prior Buenrostro Universal primer that has been modified to introduce an i5 index sequence (n)_8_ at the appropriate location based on Illumina primer designs. The ATAC UDI Index Read 1 primer corresponds to the indexed Buenrostro primer designs, although we did not retain the same i7 barcodes. Because not all index combinations are compatible with one another, we used index pair combinations reported for the New England Biolabs NEBNext Multiplex Oligos for Illumina (96 Unique Dual Index Pairs) set (catalog number E6440, New England Biolabs, Beverly, Massachusetts), replacing the (n)_8_ sequences above with the appropriate i5 and i7 barcode. Primers were synthesized with a terminal phosphorothioate bond (*) by IDT (Integrated DNA Technologies, Coralville, Iowa). No major differences were observed with library amplification or sequencing efficiencies between the UDI primer sets and the original Buenrostro designs.

Amplified libraries were checked on an Agilent Bioanalyzer (Agilent Biotechnologies, Santa Clara, California) to confirm the presence of distinct open and nucleosome-associated fragment sizes. Twelve embryos total were processed for the comparison between wild-type and ‘patternless’ mutant embryos: three embryo biological replicates per timepoint (NC14+12’ or +72’) and per genotype. For ATAC on *opa* mutant embryos, 22 embryos were collected from a cross between *opa/+* heterozygous adults and sequenced in two separate library pools. This yielded five wild-type embryos, three *opa* mutant embryos, and 14 heterozygous embryos. For ATAC-seq on NC13+12’ wild-type and *tub>opa*/+ embryos, six embryos were collected per genotype.

### ChIP-seq

Per replicate, 200 cellular blastoderm stage *w^1118^* or *opa-3xmyc* embryos were collected, crosslinked and ChIPped as described in [72] for a total of three independent biological replicates. The ChIP antibody was an anti-myc tag polyclonal antibody from Sigma-Aldrich (C3956, Sigma-Aldrich, St. Louis, Missouri). ChIP-seq libraries were prepared using the NEBNext Ultra II library prep kit with the NEB Unique Dual Index primer set kit (New England Biolabs). Samples were sequenced at the Northwestern University NuSeq Core Facility on an Illumina HiSeq 4000 on single-end 50 bp mode. Demultiplexed reads were run through TrimGalore! (https://github.com/FelixKrueger/TrimGalore) and mapped to the dm6 assembly of the Drosophila genome using Bowtie2 with options -fivePrimeTrim 5 -N 0 –local [102]. Suspected optical and PCR duplicates were marked by Picard MarkDuplicates (http://broadinstitute.github.io/picard/). To determine a set of high-confidence peaks, we called peaks using relaxed stringency with MACS2 [75] followed by determination of reproducible peak regions per replicate using the Irreproducible Discovery Rate algorithm [76]. First, we used Bedtools bamtobed (https://bedtools.readthedocs.io/en/latest/) to convert .bam formatted mapped, filtered primary sequencing reads to .bed format, which facilitated generation of pseudoreplicate datasets. Pseudoreplicates were generated by randomly splitting each replicate sample into two .bed files. Overall pseudoreplicates were generated by pooling all replicates for either *w* or *opa-3xmyc* and randomly splitting reads into two separate .bed files. Several peaks files were generated using MACS2 in preparation for IDR analysis. First, peaks for each individual *opa-3xmyc* sample (n = 3) were called against the pooled *w* sample data as a control. Having previously determined the MACS2 parameter *-d*, we bypassed model building (--nomodel) and manually specified the expected fragment size (--extsize 149). Relaxed conditions were specified by designating the option -p 1e-3 as recommended by IDR. Samples were scaled to the larger dataset (--scale-to large). A second peaks list (n = 1) was generated for the pooled *opa-3xmyc* samples against the pooled *w* samples using the same MACS2 options. A third set of peaks (n = 6) were called for each pseudoreplicate for each biological replicate of *opa-3xmyc* against the pooled *w* samples, using the same MACS2 options. Finally, a fourth set of peaks (n = 2) were called for the two pseudoreplicates of the pooled *opa-3xmyc* data against the pooled *w* data, using the same MACS2 parameters. We then performed IDR analysis on this set of peaks with threshold values of 0.02 for individual replicates and 0.01 for the pooled data sets. IDR was performed for pairwise comparisons between replicates (e.g., rep 1 vs rep 2, rep 2 vs rep 3, and rep 3 vs rep 1) using as a reference the peak list from pooled *opa-3xmyc* samples. IDR options were --input-file-type narrowPeak --rank p.value --soft-idr-threshold 0.02 and --use-best-multisummit-idr. IDR was then subsequently performed on each pair of pseudoreplicates for each individual biological replicate (e.g., rep1 pseudo 1 vs rep 1 pseudo 2…) using the same IDR options. Finally, IDR was performed on the pooled sample pseudoreplicates using the same IDR options except --soft-idr-threshold was 0.01 instead of 0.02, per the recommendation of the IDR instructions. The largest number of reproducible peaks between replicates was 881, and so we took the top 881 peak regions (ranked by p-value) from the list of pooled peaks and filtered out two regions that mapped to non-canonical chromosomes, leaving 879 high confidence peaks. Consequently, this is a conservative estimate of true Opa binding sites but is determined by more rigorous criteria (reproducibility between biological replicates) than either arbitrary p-value thresholding or simply taking the top N peaks.

### ATAC-seq Analysis

Demultiplexed reads were trimmed of adapters using TrimGalore! and mapped to the dm6 assembly of the Drosophila genome using Bowtie2 with option -X 2000. Suspected optical and PCR duplicates were marked by Picard MarkDuplicates. Mapped, trimmed, duplicate marked reads were imported into R using the GenomicAlignments [103] and Rsamtools libraries (http://bioconductor.org/packages/release/bioc/html/Rsamtools.html), filtering for properly paired, non-secondary, mapped reads, with map quality scores greater than or equal to 10. Reads with mapped length less than or equal to 100 bp were considered to have originated from ‘open’ chromatin.

For the *opa* mutant ATAC-seq experiment, embryos were collected blind to the zygotic genotype, but because experiments were performed on single embryos, genotypes could be determined post-hoc by calling SNPs over the *opa* locus using BCFtools [104].

Peak regions for all accessible regions present between ZGA and gastrulation were called using MACS2 with options -f BAMPE -q 1e-5 --nomodel on a merged dataset comprising all ‘open’ ATAC reads from both wild-type and ‘patternless’ mutant embryos, and both NC14+12 and +72 timepoints.

DEseq2 [47] analysis was performed by counting the number of ‘open’ ATAC seq reads overlapping each peak identified by MACS2. For the comparison between wild-type and ‘patternless’ mutant embryos over NC14+12 and +72, the design parameter passed to DESeq2 was ∼genotype + time + genotype:time. For other comparisons (*opa* gain or loss of function at single timepoints) the design was ∼genotype only. Regions with an adjusted p-value of less than 0.01 and absolute log_2_ fold-changes > 1 were considered to be statistically significant in the relevant test.

To determine the different dynamic peak classes (Figure 4), we first undertook a clustering based approach to explore how many different classes we could identify within the dataset. To do this, we first averaged the number of reads per peak region for each sample and then scaled the data for each peak region by dividing the mean reads for a peak region by the sum of all mean reads for a peak region (i.e., so that the sum of the scaled reads for a peak region, sum_region_(wt 12, wt, 72, mut 12, mut 72), would equal 1). Next, we performed k-means clustering with a variable number of cluster centers, and plotted the average per-cluster profile to visualize average behavior of clustered regions. We found that 10 was the minimum number of cluster centers that would capture all the unique patterns present in the data, that fewer clusters would combine similar but qualitatively different classes, and more clusters would subdivide clusters into relatively similar subsets. We note that over the two-year course of this study, attempts were made to replicate this analysis on three different Apple (Apple Computer Inc., Cupertino, California) computers running different base operating systems, versions of R, and versions of dependent libraries. For reasons that are not clear, clustering based approaches were not strictly reproducible across differently configured computers despite identical input datasets and identical scripting of the analysis code. To be clear, similar cluster types, and minimum cluster number were called across different systems, however the numeric order of the clusters, as well as the number of peaks assigned to each cluster would vary between systems. Because of this, we could not rely on clustering alone to reproducibly describe our results. Therefore, we took an alternative approach to categorizing peak classes that depended on the statistical output of DESeq2, whose output was identical across different computer systems. We paired statistical tests from DESeq2 to define classes. For instance, to define regions that were uniformly open early and uniformly lost accessibility by gastrulation, we required both wild-type and mutant samples to have a statistically significant difference across timepoints, with a log2 fold change of -1 or less. On the basis of these paired DESeq2 criteria, we reproduced each of the 10 peak class types predicted by clustering. We note that the final number of categorized peaks (2917) is substantially lower than the number of peaks that score above the significance + log2 fold change thresholds of 0.01 + |1| (6775). This is due to the fact that we require pairs of DESeq2 tests to score above significance thresholds for assignment to a class. The remaining 3858 regions only score as significantly different in one of the four tests.

### Generating reporter constructs based on ATAC-seq data

We used ATAC-seq coverage to delineate the sequence to test for potential enhancer activities associated with *wg-2.5*, *wg-1*, *odd-late*, *slp1 “5”*, *18w 1* and *18w 2*. In general, coverage at peak regions was plotted on the UCSC genome browser and views were zoomed out to identify flanking regions of low accessibility. We hypothesized that functional genomic elements would be defined by extended regions of accessible chromatin flanked by inactive, low-accessibility regions. Primer pairs were designed to amplify peak regions plus a small amount of flanking ‘low-accessibility’ DNA and were cloned into a Gateway entry vector pENTR (Thermo Fisher Scientific, Waltham Massachusetts). The exception to this overall strategy was in the case of the *wg-2.5* and *-1* regions which are adjacent to one another (Figure 3A). in which case we delineated the two regions by the midpoint between the two major peaks of chromatin accessibility. Once we had generated one pENTR-enhancer clone by TA-cloning of a PCR product, it was no longer necessary, efficient, or desirable to perform TA cloning. Subsequent pENTR clones were made by excising the original insert and replacing with new candidate enhancer fragments via Gibson Assembly (NEB HiFi Assembly Kit, New England Biolabs). The candidate enhancer fragments were shuttled to the transgenic vector pBPGUw [105] upstream of a minimal synthetic promoter sequence driving Gal4 using standard Gateway cloning techniques (LR Clonase, Thermo Fisher Scientific). Transgenic lines were established as described above by insertion into the *attP40* landing site.

The genomic coordinates corresponding to regulatory elements tested in this study are: *odd-late*: chr2L:3602087-3603534; *slp1 “5” variant*: chr2L:3817360-3818884; *wg -1:* chr2L:7305311-7306265; *wg -2.5*: chr2L:7303832-7305569; *18w 1*: chr2R:20091666-20092774; *18w 2*: chr2R:20071902+20073306.

### In situ hybridizations

Probe sequences for *in situ* hybridizations were generated by PCR against cDNA clones for the target transcript. Reverse primers included a leading T7 promoter sequence to facilitate probe synthesis by *in vitro* transcription using an Ambion MegaScript T7 kit (Thermo Fisher Scientific) supplemented with digoxigenin-11-UTP (Roche, Sigma-Aldrich) following standard procedures. *In situ* hybridizations were performed according to standard procedures using a hybridization temperature of 65°C, which was found to significantly reduce background compared with 56°C. For reducing bias in the evaluation of gene expression in ‘patternless’ mutant embryos, color development reactions were performed side-by-side with a wild-type control. Color development was evaluated by observing only the wild-type sample and stopping both reactions once it was determined that wild-type samples had just reached optimal signal. Staining levels in mutant embryos were then evaluated following termination of the color development reaction. For evaluating sensitivity of a reporter to *opa* loss of function, we performed *in situ* hybridizations against *gal4* on two samples in parallel, a wild-type sample of *test enhancer>gal4* homozygous males crossed to *w^1118^* females, and embryos from a cross between *opa^IIP32^/ TM3, Sb twi>gal4 UAS>GFP* females and *w*; *(test enhancer>gal4)*; *opa^IIP32^/ TM3, Sb twi>gal4 UAS>GFP* males, which allowed us to distinguish test enhancer>reporter expression in *opa* homozygous mutant embryos by their lack of *twi>gal4* staining in the ventral mesoderm. Color development was monitored in the wild-type sample, and mutant embryos were ultimately hand selected from the *twi>gal4*-marked sample, after ensuring that wild-type expression patterns matched the *twi>gal4*-positive expression patterns, save for the broad domain of *twi>gal4* expression. Images shown in figure panels are of similarly staged wild-type samples and *twi>gal4*-negative (*opa* mutant) samples imaged under identical conditions. T7-antisense Dig-UTP riboprobe template-generating primer sequences are as follows:

*gal4-F*: 5’-TGCGATATTTGCCGACTTA-3’

*gal4-R*: 5’-TAATACGACTCACTATAGGGAACATCCCTGTAGTGATTCCA-3’

*ems-F*: 5’-AGCATCGAGTCCATTGTGGG-3’

*ems-R*: 5’-TAATACGACTCACTATAGGGTATCTCCTGGCCGCTTCTCT-3’

*Kr-F*: 5’-CTGGATGTGTCGGTGTCTCC-3’

*Kr-R*: 5’-TAATACGACTCACTATAGGGCGGTAACAAGCATACGGT-3’

*byn-F*: 5’-ACGAACCACGTGTGCATCT-3’

*byn-R*: 5’-TAATACGACTCACTATAGGGTAGGAACTGCTGAAGCAACCA-3’

*ush-F*: 5’-GAAGGTGCGAGTGAAGTGGA-3’

*ush-R*: 5’-TAATACGACTCACTATAGGGCGAATGGGCTTGTTTTT-3’

*sog-F*: 5’-CCATTGTGTGTGCCAGTGTG-3’

*sog-R*: 5’-TAATACGACTCACTATAGGGCACTTGGTCTCTCCGTTCA-3’

*sna-F*: 5’-CCGGAACCGAAACGTGACTA-3’

*sna-R*: 5’-TAATACGACTCACTATAGGGCCAGCGGAATGTGAGTTTGC-3’

## Data Availability

Data will be made publicly available upon publication of the peer-reviewed manuscript. Data will be made available to reviewers during the peer review process.

## Acknowledgments

We gratefully acknowledge Eric Wieschaus, in whose laboratory this project was initiated, for his generosity, support, and encouragement. We also thank Angelike Stathopoulos, Melissa Harrison, Urs Schmidt-Ott, Jeff Farrell, Hernan Garcia, Jacques Bothma, Eileen Furlong, Nicolas Gompel, Dan McKay, Carole LaBonne, Rich Carthew, Erik Andersen, and Greg Beitel for helpful conversations. This work was supported by a Searle Leadership Fund for Life Sciences Award and Northwestern University Startup Funds to S.A.B.

